# Pinhole Optical Tweezers: Extending the Photobleaching Lifetime in the Presence of an Optical Trap by Wavefront Engineering

**DOI:** 10.1101/808352

**Authors:** Zheng Zhang, Joshua N. Milstein

## Abstract

We present a new method for combining optical tweezers with single-molecule fluorescence in an engineered geometry we have coined a ‘pinhole’ optical trap. By utilizing an appropriately constructed Laguerre-Gaussian (LG) or ‘donut’ beam, and applying force along the axis of the trapping laser, one can maintain a low-intensity region of near-infrared (IR) light directly below the optical trap in which a biomolecule may be probed by both force spectroscopy and fluorescence. We show that within this region of low IR light intensity, the photobleaching lifetime of Alexa-647, an organic dye that is particularly sensitive to the high intensity trap light, can be significantly extended. This approach enables us to spatially separate the trap light from the fluorescence illumination without the need to physically separate, by many micrometers, the optical trap from the biological sample.

## Introduction

Optical tweezers and single-molecule fluorescence are two powerful biophysical techniques that often provide complementary information on both the static and dynamic features of a macromolecular system. Force spectroscopy can probe a range of biomechanical processes, from the energy landscapes of proteins and nucleic acids to the load-dependent kinetics of enzymatic molecular machines^1–3^. Single-molecule fluorescence measurements, on the other hand, can yield direct information on chemical kinetics such as explicitly identifying residues involved in local, structural changes to a protein^4, 5^. Employing both techniques in tandem can provide significant new insight into molecular mechanisms inaccessible, or at least less accessible, to either technique alone.

Combining these two techniques, however, is challenging. The primary obstacle is that the high intensity light from the trapping laser tends to drastically increase the bleaching rate of fluorescent dyes through multi-photon processes. Though the mechanism is not well understood, previous work suggests that the enhanced photobleaching is caused by the absorption of a near-infrared photon while in the lowest, excited singlet state *S*_1_. This event drives the dye into a higher order singlet state, from which intersystem crossings may occur that readily form nonfluorescent dye cations^6^. This is in contrast to the traditional route whereby a dye excited to *S*_1_ crosses into a triplet state where it may generate reactive oxygen species (ROS) that degrade the fluorophores^7^. Although some fluorophores, such as TMR^6, 8^, are not significantly affected, many popular dyes suffer from large increases to their bleaching rates. In particular, red-absorbing dyes, such as Alexa-647, show the most drastic reduction in their lifetimes in the presence of an optical trap^6^.

This enhancement of photobleaching is only significant near the focus of the trapping laser, so it can be avoided altogether by spatially separating the dyes from the optical trap^9–12^. However, this approach typically requires the use of long DNA linkers, which can contribute a significant level of thermal noise to a measurement and severely restricts the experimental geometry^13, 14^. An alternative approach is to interlace the trapping and excitation beams in time such that they are temporally separated^15, 16^. By ensuring both lasers are not incident on the dye simultaneously, the increase in photobleaching can not only be eliminated, but native photobleaching may even be reduced^17^. While an effective solution, this approach is technically challenging, so has not gained acceptance beyond a very limited number of labs.

Here we introduce a new method that utilizes wavefront engineering to combine single-molecule fluorescence with axial optical tweezers^18^. By performing axial force spectroscopy with a carefully constructed Laguerre-Gausssian (LG) donut beam, a conical, ‘dark’ region of significantly lower intensity, infrared light is generated directly below the trapped microsphere. Since extensions along the axis of the trap laser essentially maintain the position of this dark region, the trap light and fluorescence illumination are separated without the need for long DNA linkers. We refer to this device as a ‘pinhole’ optical tweezers, and have found that the bleaching rate of surface mounted Alexa-647 dyes, which are highly sensitive to the near-IR trap light, is significantly reduced when illuminated within the dark region of such a device.

## Results

### Wavefront Engineering

For our optical tweezers system, we both manipulate and engineer the optical trap with the use of a phase-only spatial light modulator (SLM). Coarse positioning of the trap within the sample is achieved by applying the phase hologram:

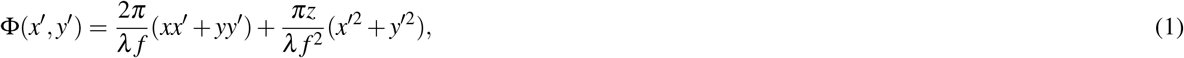

where (*x*′, *y*′) are the SLM pixel coordinates and *f* is the focal length of the objective. The two terms of Eq. 1, in order, are a blazed grating to displace the trap laterally in *x* and *y*, and a Fresnel lens for axial displacements along *z*.

The simplest way to generate a Laguerre-Gaussian donut mode (i.e., 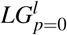) is to apply an additional phase wrap to the hologram Φ_*LG*_ = *mφ*, where *φ* is the polar angle on the SLM and *m* = ± *l* is an integer denoting the azimuthal index^19, 20^. This approach, however, does not produce a pure LG donut mode, rather it generates a superposition of higher-order radial *p* modes for a given azimuthal mode *l*. The result is a series of concentric, annular rings, albeit with the majority of the power in the *p* = 0 mode. Note, there exist more advanced, holographic methods for generating purer 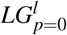 donut modes; however, these approaches all lead to a significant loss of power (*>* 50%), which translates into a much weaker optical trap^21^. We have found that the approximate donut modes generated by a simple phase wrap are sufficient for our purposes.

Another necessary factor to consider when generating an LG donut mode trap is the polarization of the light. Tight focusing of a donut beam though a high-NA objective can lead to variations in the intensity and phase profile near the focus. We had previously shown that these polarization effects can alter the strength of an optical trap created with an LG donut mode^22^. In fact, to create a true dark region within the centre of the annular trap light, it was necessary to circularly polarize the trap light and to match the handedness of the polarization to the phase wrap of the computer generated hologram. With linearly polarized light, the trap axis began to fill with IR light, and the central dark region almost vanished when trapping with circularly polarized light that spirals opposite the phase of the hologram.

In view of our previous findings, we employ vectorial Debye theory^23, 24^ to discern the importance of polarization effects as one moves away from the focus; to ask if they affect the intensity within the dark region at the coverslip surface where we would like to perform fluorescence measurements. The results of our numerical simulations are presented in Fig. 1. While the intensity along the central axis of the trapping laser is strongly affected by the relative orientation of the laser polarization and phase wrap near the focus, the importance of aligning the two greatly diminishes as one moves away from the focus. In fact, at a little over 1 *µ*m below the focus the difference in the central intensity between an aligned and anti-aligned beam becomes negligible. These results suggest that polarization effects are not critical in creating a dark region at the coverslip unless one is trapping extremely close to the surface.

**Figure 1.**
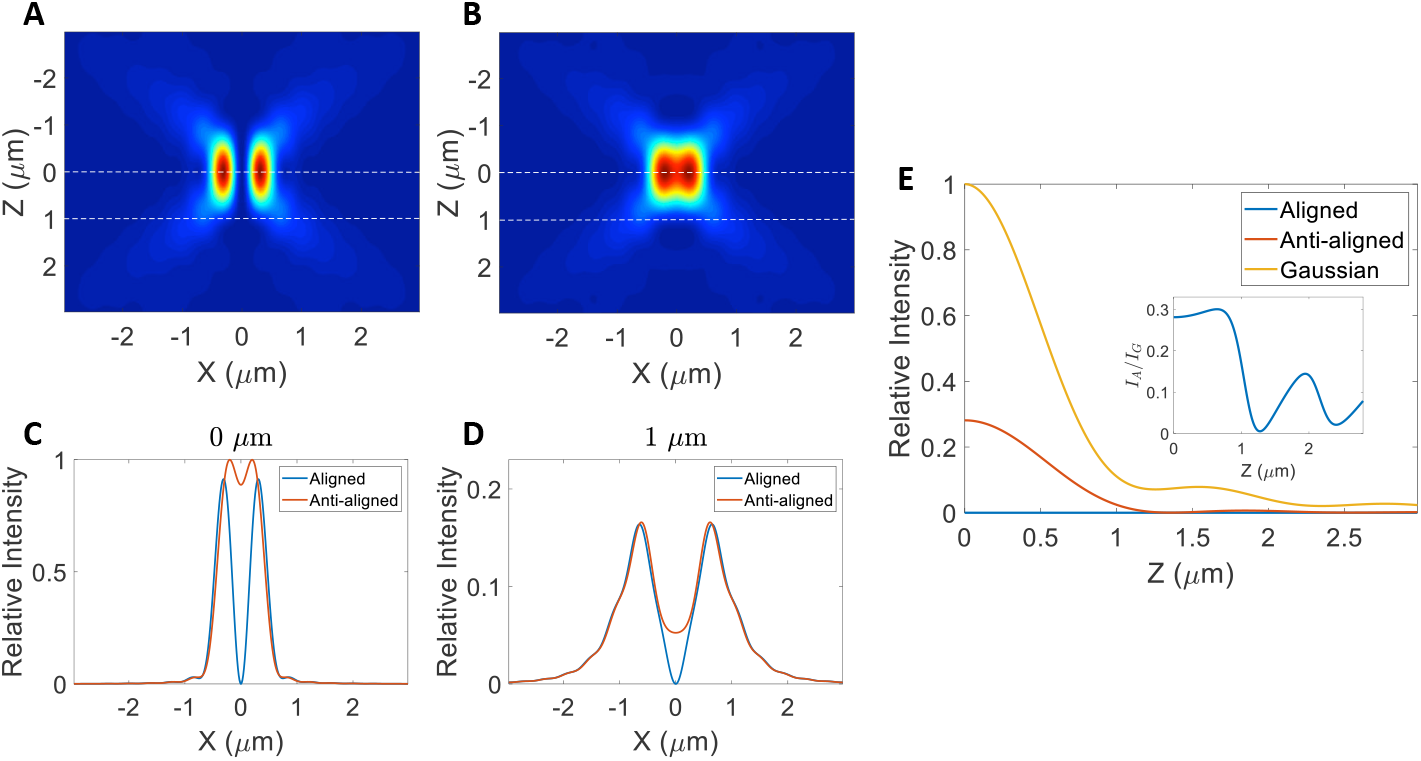
Numerical simulations of a tightly focused *l* = 1 donut beam. Cross sections of the intensity profile for an A) aligned and B) anti-aligned donut beam. Relative intensity profile of an aligned and anti-aligned donut beam C) at the focus and D) 1 *µ*m below the focus. These locations are indicated in A) and B) by the dashed lines. E) Comparison of the relative intensity of an aligned, anti-aligned and Gaussian beam along the trap axis (*X* = 0). Insert: ratio of intensities between the anti-aligned (*I*_*A*_) and gaussian (*I*_*G*_) beam.

### Photobleaching Measurements

We next present a series of measurements showing that the photobleaching lifetime of an organic dye can be extended in the presence of a donut beam optical trap within the dark region below the trap. Figure 2A illustrates the experimental configuration. A coverslip coated with Alexa-647 dyes is carefully positioned below an optical trap. This is conveniently achieved by considering the spot size that results from imaging surface bound dyes in the presence of a Gaussian trap. By scanning the trap up and down with the SLM, so as to minimize the spot size, the trap focus can be placed at the coverslip surface and can then be displaced to a precalibrated distance above the surface. Epifluorescence images of the surface under continuous exposure by a 638 nm excitation laser are then collected. In the presence of a Gaussian trapping beam, a dark hole quickly appears due to accelerated photobleaching induced by the high intensity trap light (Fig 2B top). However, active dyes remain visible after several minutes within the center of the partially bleached region just below the focus of a donut beam (Fig 2B bottom).

**Figure 2.**
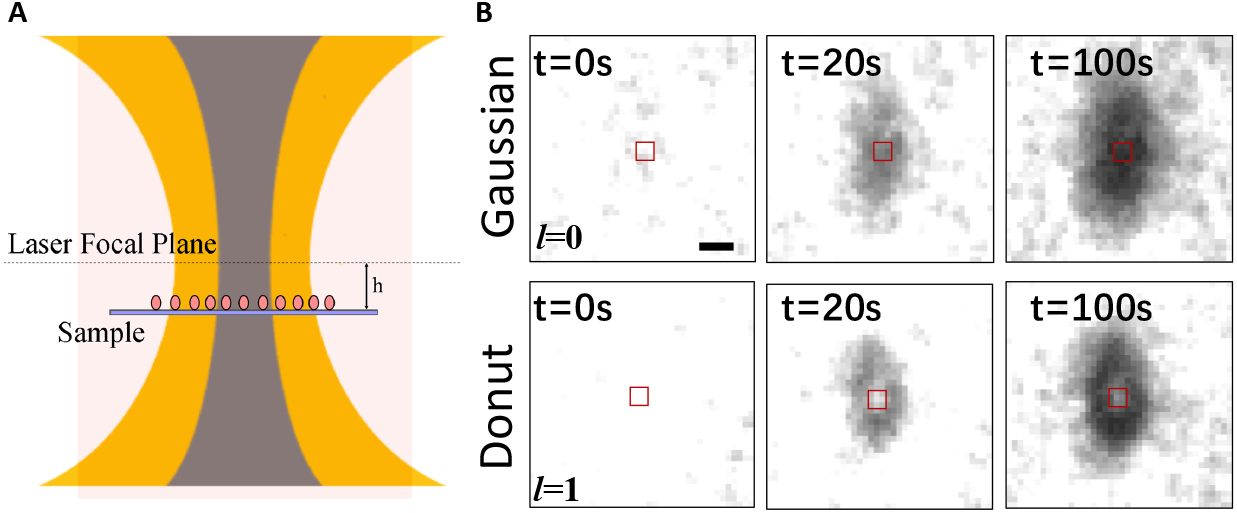
A) Illustration of experimental imaging setup. B) Epifluorescence images of Alexa-647 coated coverslips in an imaging buffer containing PCA/PCD. From left to right (t = 0s, 20s and 100s), the images show spatial variation in the enhancement of photobleaching due to a Gaussian (top) and Donut (bottom) beam optical trap positioned *h* = 1 *µ*m above the coverslip surface. The red squares indicate the measured 3×3 pixel region. Scale bar equals 1 *µ*m (∼ 6 pixels).

These results are quantified in Fig. 3. For this analysis, we select a 3×3 pixel region centred directly below the trap and sum the pixel intensities. This measurement is repeated multiple times and averaged together to yield the curves in Fig. 3 (A,B,D and E), which are then fit with a double exponential of the form *f* (*t*) = *A* exp(−*t/τ*_1_)+ *B* exp(−*t/τ*_2_)+*C*. The effective photobleaching lifetime is defined from the fits as *τ* = (*Aτ*_1_ + *Bτ*_2_)/(*A* + *B*), which can also be thought of as a normalized measure of the total photon output of the dyes. We note that an analysis of a 2×2 pixel region yields similar values of *τ*, but requires significantly more averaging (i.e., more measurements), while analyses of larger pixel regions are increasingly affected by inhomogeneities in the trap illumination.

**Figure 3.**
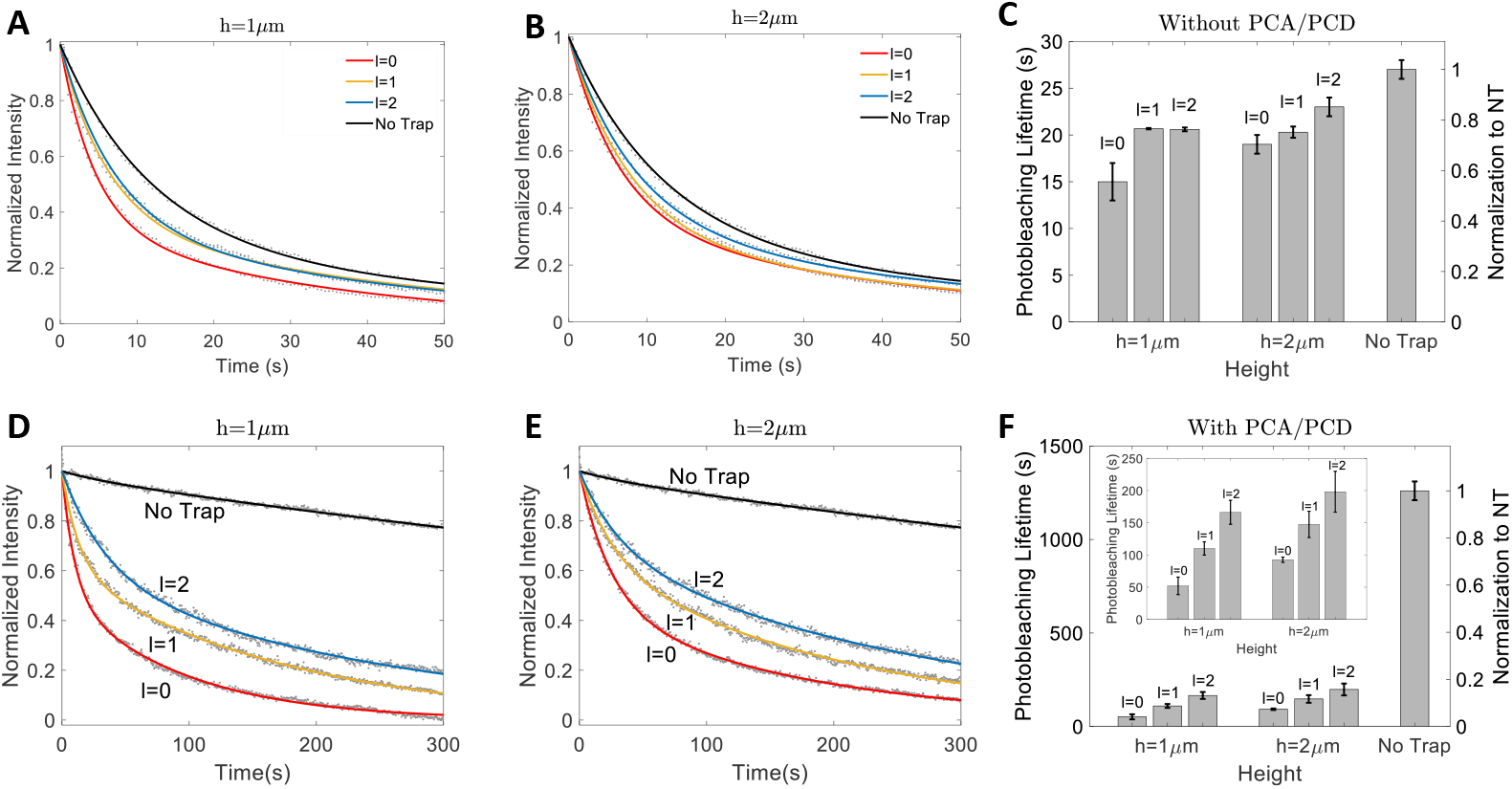
Comparison of photobleaching enhancement within a 3×3 pixel region centered below the optical trap. Intensity decay curves in PBS buffer at A) 1 *µ*m and B) 2 *µ*m below the trap for no trap, an *l* = 1 and *l* = 2 donut beam, and a Gaussian beam. C) Absolute and normalized photobleaching lifetimes extracted from the data. D-F) same as A-C, respectively, in PBS with the oxygen scavenging system PCA/PCD. The insert to F is an enlarged view of the absolute lifetimes.

Figures 3A and 3B show the intensity decay curves of Alexa-647 at 1 *µ*m and 2 *µ*m below the trap when there is no oxygen scavenging system in the imaging buffer. In this case, even without the trap light, the dyes quickly photobleach in well under a minute. At 1 *µ*m below the focus, the photobleaching lifetime is a little more than half the lifetime in the absence of the trap light. For an *l* = 1 donut beam, we observe an approximately 40% improvement in the absolute lifetime, with little or no additional gains resolvable in our measurements for an *l* = 2 beam. Positioning the trap 2 *µ*m above the coverslip yields less of an improvement, although the gains are now clearer for an *l* = 2 beam. Note that placing the trap at 2 *µ*m also increases the photobleaching lifetime in the presence of a Guassian beam by reducing the trap light intensity at the surface (compared to when it’s positioned at 1 *µ*m).

The absolute gains are even more pronounced in the presence of the PCA/PCD oxygen scavenging system (Figs. 3D-F). For a trap placed 1 *µ*m above the surface, compared to a Gaussian beam of the same intensity, the photobleaching lifetime can be increased by ~ 2x for an *l* = 1 donut beam and ~ 3x for an *l* = 2 donut beam (with slightly less improvement in the photobleaching lifetime measured at 2 *µ*m). We note that while these are significant gains in terms of the absolute photobleaching lifetime, Alexa-647 is extremely stable in the presence of PCA/PCD when their is no trap light present. The longest photobleaching lifetimes we measured below a donut beam optical trap are still only a fraction of the lifetimes measured without the trap.

We also considered the photobleaching lifetime as a function of the trap power at the back focal plane of the objective. Here we only considered the case of positioning the optical trap 1 *µ*m above the surface. In our measurements, we typically see a monotonic decrease in the photobleaching lifetime as the trap power is increased from 50 mW - 220 mW (Fig. 4). This trend is in agreement with previous reports on the photobleaching lifetime in the presence of an optical trap^6, 17^. Moreover, these results show that significant gains can be made in increasing the photobleaching lifetime by working with a donut beam over a Gaussian trap, in particular, at high powers (which translates to higher optical trap strengths).

**Figure 4.**
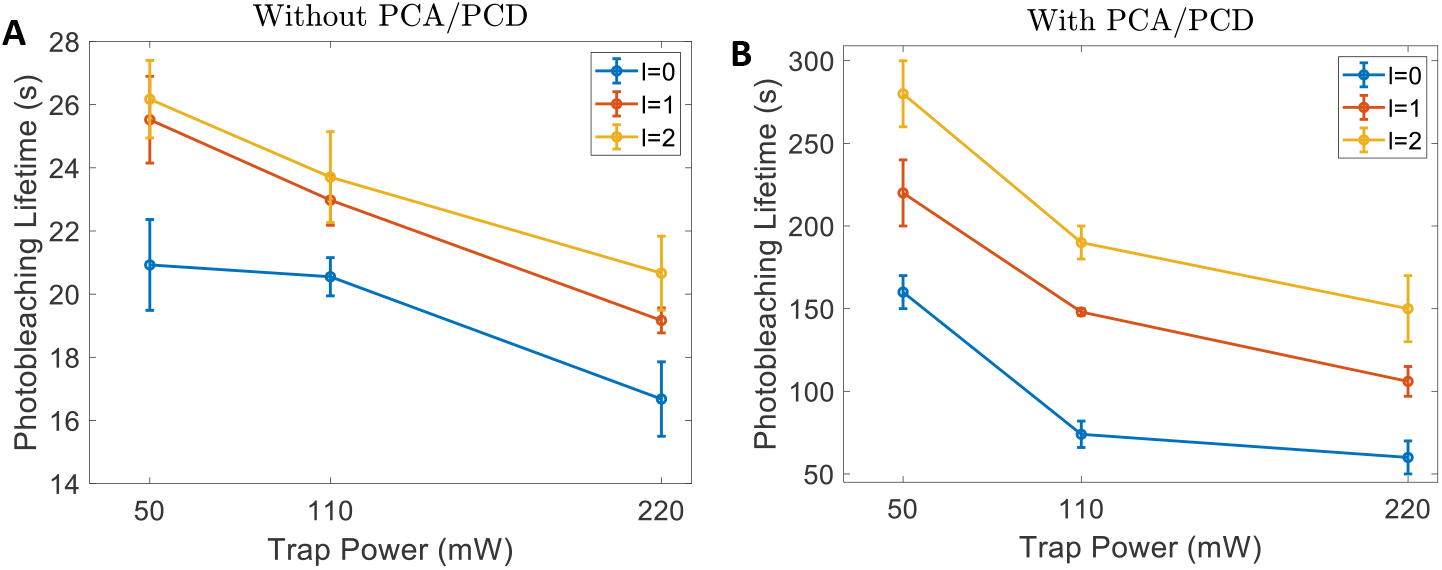
Photobleaching lifetimes as a function of the trap laser intensity A) in PBS and B) in PBS with the oxygen scavenging system PCA/PCD. For these measurements, the optical trap was positioned 1 *µ*m above the coverslip surface.

## Discussion

Our measurement and quantification of an effective photobleaching lifetime showed that a pinhole optical tweezers can significantly increase the lifetime of a surface bound organic dye when performing simultaneous optical trapping and fluorescence experiments. These results, however, still underestimate the improvement in photobleaching lifetimes resulting from this trapping geometry. To understand why, let us reconsider our previous analysis in light of Fig. 5A, which illustrates a complication to interpreting our measurements. The intensity of dyes within a relatively small region, centred along the axis of the optical trapping laser, are measured as a function of time. However, the point spread function (PSF) originating from a single dye has a FWHM of ~ 580 nm, which can extend over approximately 4 pixels. This means that the PSF of dyes outside of the central region will overlap with this region contributing to the total measured intensity. In fact, if the area of this central region is relatively small, and we assume the dyes are randomly dispersed on the coverslip with a roughly uniform density, the majority of the light intensity within the central region results from dyes outside its perimeter. In our measurement, approximately 70% of the light measured within the considered 3×3 pixel region (*R* ~ 240 nm) originates from outside dyes overlapping with the region of interest (Fig. 5B).

**Figure 5.**
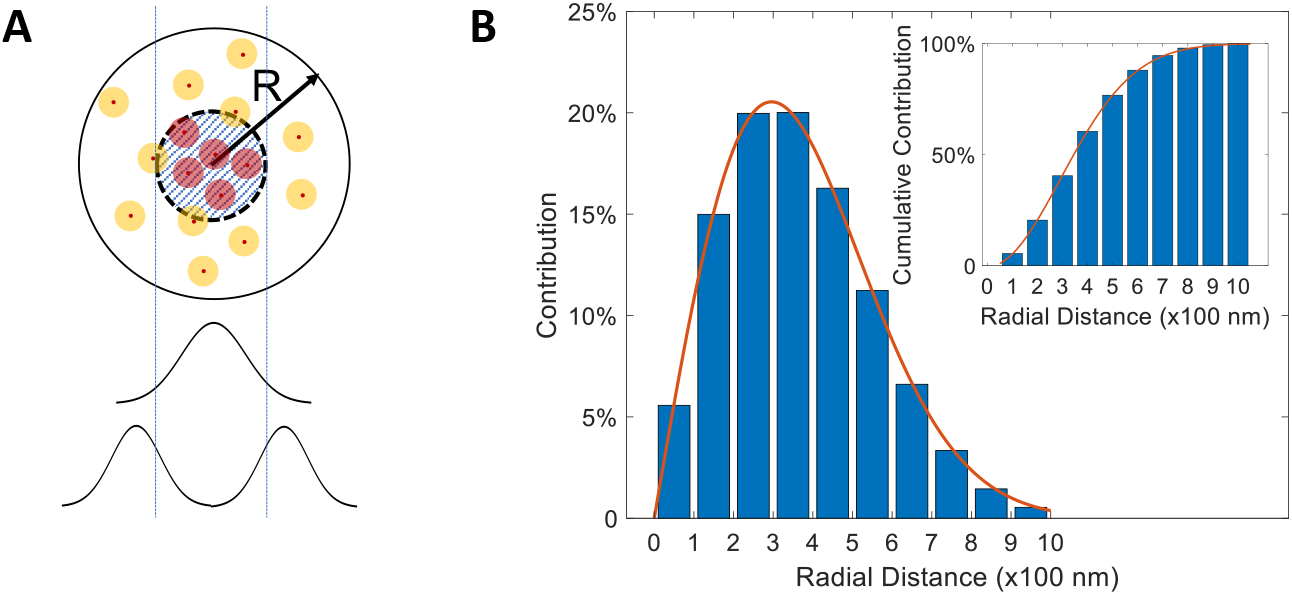
A) The intensity of the central region (shaded) arises from the PSF of dyes within the region (red circles) and overlap of the PSF of dyes outside the region (orange circles). A Gaussian beam accelerates photobleaching in the central region while a donut beam accelerates photobleaching around the central region. For clarity, this image is not to scale. B) Percent contribution to the central intensity as a function of radial distance (*R*). The solid line is the analytic function *f* (*x*) = *ax* exp(*−x*^2^/2*b*^2^), with normalization constant *a* and *b* = 296 nm corresponding to the width of the dye PSF. Insert: cumulative contribution to the central intensity. The line indicates (1 − *f* (*x*)) × 100%.

These exterior, overlapping PSFs result in a background signal that has a temporal dependence. Let us first consider our measurements in PCA/PCD containing buffer. Here, the background (i.e., no trap) photobleaching rate is so much slower than any of the other photobleaching processes that, as a first approximation, we may assume it to be negligible. We next differentiate two dynamic processes that contribute to our measurements: 1) the rapid photobleaching of dyes with rate 1/*τ*_*f*_ in the presence of high intensity trap light and 2) the slower photobleaching of dyes with rate 1/*τ*_*s*_ in low intensity regions of the trap beam. Consider illumination by a Gaussian beam. In this case, our double exponential fit can be interpreted as measuring the photobleaching of dyes within the central region by the fast time constant *τ*_*f*_, with the slower time constant *τ*_*s*_ quantifying the decay of external dyes that overlap with the central region (Fig. 5A). For a donut beam, however, the exponential fit needs to be reinterpreted. In this case, the slow time constant *τ*_*s*_ quantifies the intensity decay of dyes within the central region, while the fast time constant *τ*_*f*_ accounts for photobleaching of external, overlapping dyes (Fig. 5A). We corroborated this reasoning, by measuring a 3×3 pixel region off centre and on the intensity maximum of the donut beam. As predicted, we found the fast time constant to be within the experimental error of that measured within the central region. In Fig. 6, we summarize our reevaluated results for the PCA/PCD buffer measurements. Compared to a Gaussian trap, we see that a pinhole optical tweezers can actually extend the useful lifetime of Alexa-647 by over an order-of-magnitude within the dark region of the trap. For our measurements in PBS without PCA/PCD, the background photobleaching rate of Alexa-647 cannot be neglected. This prevents us from interpreting the double exponential fits to our measurements in the simple way outlined above for the PCA/PCD case. However, it’s fair to say that the results shown in Figs. 3A-C should be considered as lower bounds on the extension of the photobleaching lifetime in the presence of a pinhole tweezers.

**Figure 6.**
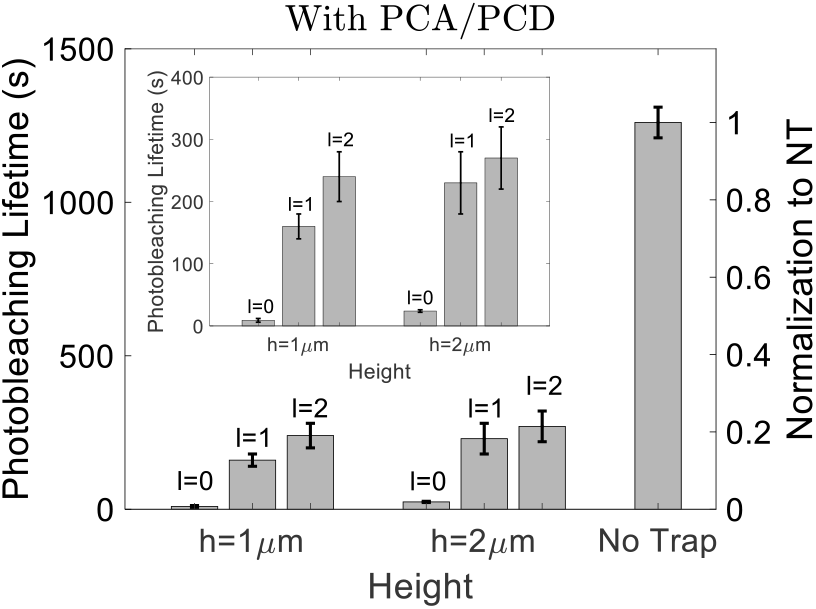
Absolute and normalized photobleaching lifetimes, in PBS with the oxygen scavenging system PCA/PCD, as quantified, according to our model, by a single-time constant. Insert: enlarged view of the absolute lifetimes.

We have shown that by optical trapping with a donut beam, an IR dark region can be maintained directly below the optical tweezers. The photobleaching lifetime of an organic dye within this dark region can be significantly extended (by over an order-of-magnitude) from what it would be in the presence of a conventional optical tweezers that employs a Gaussian beam. And if force extension measurements are performed along the axis of the trap laser, the geometry will preserve the position of this dark region. We coined this technique an axial pinhole optical tweezers, which can serve as a simplified geometry in which to perform dual force spectroscopy and single-molecule fluorescence measurements. The resulting method should be a welcome addition to the toolbox of single-molecule techniques.

## Methods

### Optical Tweezers Setup

The holographic tweezers portion of the setup was initially described in^22^. The original setup, extended to include fluorescence imaging, is presented in Fig. 7. Light from a 1064 nm Nd:YAG laser (4 W, Coherent BL-106C) is optically isolated and linearly polarized by a half wave plate (HWP) in combination with a polarising beam splitter (PBS), which also allows for manual tuning of the laser power. The light is then passed through a pinhole to yield a highly symmetric, circular beam, which is then reflected off a phase only spatial light modulator (SLM) (Hammamatsu X10468) at a highly oblique angle and 4-f imaged onto the back focal plane (BFP) of a high-NA, oil-immersion objective (Olympus PlanApo 100x, 1.4 NA). An iris placed in an intermediate Fourier plane blocks the zeroth order (unmodulated) and any unwanted higher-orders of light reflected off the SLM (i.e., only the first order light is allowed to pass). The light is then circularly polarized with a quarter wave plate (QWP) before focusing into the sample. A condenser (Olympus LUCPLanFL 40x, 0.75 NA) collects the scattered light and images it onto a position sensitive diode (PSD) (FirstSensor DL100-7-PCBA3) for back focal plane interferometry (BFPI). Positioning of the trap is controlled by the SLM, as discussed in the text, but finer nanometer scale displacements can be achieved by instead adjusting the position of the sample, which rests upon an xyz-piezo stage (Mad City Labs). For epi-fluorescence imaging, a 638nm laser is first circularized through a pinhole and focused onto the BFP of the objective. The profile of the excitation beam is a 2D-Gaussian with a 1*/e*^2^ width of ~ 24 *µ*m. In our analysis, we chose a 480 nm X 480 nm region near the centre of the Gaussian. Within this small region, the profile is essentially flat. The fluorescence signal is then isolated with a combination of filters and dichroic mirrors and imaged onto an EMCCD camera (Andor, iXion3) controlled by MicroManager^25^. The EM gain was set to 10 and the exposure time was 50 ms without PCA/PCD and 100 ms with PCA/PCD, with a 0.5 s interval between each image. All data processing was achieved by a self-written MATLAB (MathWorks, 2019a) program.

**Figure 7.**
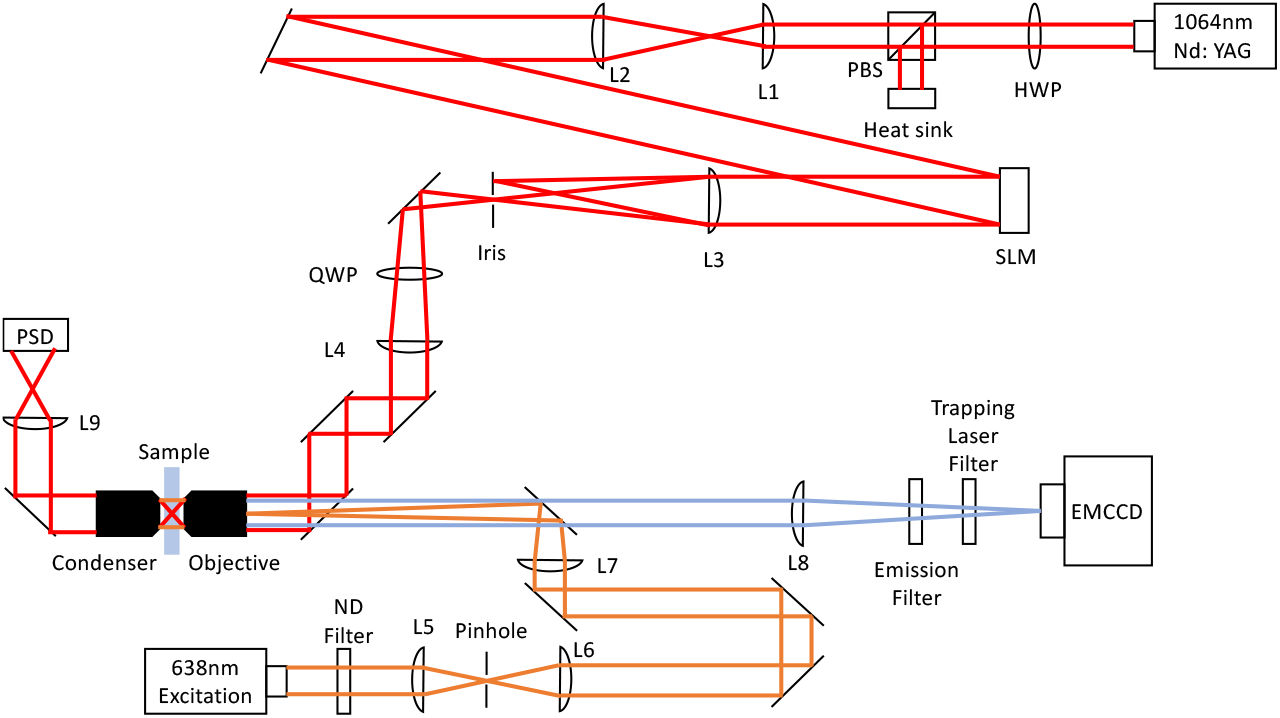
Optical configuration employed for the pinhole optical tweezers.

### Sample Preparation

Sample chambers were created by cutting a channel into a section of parafilm, which was then sandwiched between a glass slide and a coverslip, and incubated on a hot plate to adhere the layers. Two holes were initially drilled into the slide to provide access to a roughly 15 *µ*L chamber. And the coverslips were previously incubated in 0.01% (w/v) Poly-L-lysine solution (Sigma, 0.1% w/v in H_2_O, 10x dilution) overnight to facilitate dye attachment.

150 *µ*M of the organic dye Alexa-647 was diluted to 50 nM with PBS buffer. 20 *µ*L of the diluted dye solution was then pipetted into the sample chamber and incubated for 10 min to coat the dye on the coverslip surface. At this concentration, the average distance between dyes was estimated to be ~ 30 nm, so dye-dye interactions should be negligible. After incubation, the chamber was flushed 3 times with 50 *µ*L PBS (pH=7.5) buffer to flush away free dyes. For experiments that used the PCA/PCD oxygen scavenging system, an imaging buffer of 95% (volume) PBS buffer, 4% (volume) 13 mM PCA solution and 1% (volume) 50 nM PCD solution was then injected into the chamber, otherwise the dyes were simply imaged in PBS. Finally, the input ports on the chamber were sealed with tape to prevent evaporation.

## Acknowledgements

The authors thank Russell Pollari for his initial contributions to this project and for originally coining the term ‘pinhole optical tweezers’. This work was supported by a Natural Sciences and Engineering Research Council (NSERC) Discovery Grant and an Early Researcher Award from the Ontario Ministry of Research, Innovation and Science.

## Author contributions statement

Z.Z. and J.N.M conceived the experiment(s), Z.Z. conducted the experiment(s), Z.Z. and J.N.M analysed the results. All authors reviewed the manuscript.

## Additional information

All data generated or analysed during this study are included in this published article (and its Supplementary Information files). The author(s) declare no competing interests.

## Notes

#### Summary of Updates

Fixed error in the manuscript title

